# Structure of a benzosulfonamide based inhibitor – 11l - bound to a disulfide stabilised HIV-1 capsid hexamer

**DOI:** 10.1101/2023.08.16.553511

**Authors:** Michael Barnett, Lin Sun, Shujing Xu, Xinyong Liu, Peng Zhan, David C. Goldstone

## Abstract

The Human Immunodeficiency Virus Type 1 (HIV-1) continues to be a major global health issue, with infection leading to Acquired Immunodeficiency Syndrome (AIDS). Despite advances in antiviral therapy, the need for new and more effective treatments remains critical. In this study, we determined the structure of the disulfide stabilised HIV-1 capsid protein hexamer in complex with the novel antiviral capsid inhibitor compound 11l. The structure revealed the presence of six 11l molecules bound to the capsid hexamer in a conserved site shared by nuclear import factors, and other capsid inhibitor compounds such as PF74 and GS-6207. The 11l compound exhibits disorder in the benzo-sulfonamide groups and disruption in the loop between helix 8 and 9 for the capsid C-terminal domain, an important 2-fold symmetry axis for hexamer formation. Our findings provide insights into the mechanism of action of 11l as a capsid inhibitor and antiviral. These results contribute to the ongoing efforts to develop more effective antiviral treatments for the global burden of HIV and AIDS.

**Synopsis:** Insight into the mechanism of action for a novel HIV-1 capsid inhibitor antivirals are elucidated by structure of HIV-1 capsid hexamer bound to the novel inhibitor compound 11l.

## 1. Introduction

HIV-1 remains a chronic and principally incurable infection, that without medical intervention leads to Acquired Immunodeficiency Syndrome (AIDS). Treatment comes with an associated cost, with a requirement for life-long treatment and associated disease burden. Although infection cannot be readily cured, eradicating the spread of HIV-1 may be achieved provided treatment is accessible. Current treatment regimens delay the progression of the infection to AIDS and prevent the spread of HIV-1 virus, requiring a daily dose of combined anti-retroviral cocktails. Whilst effective, these treatments are costly, can have low patient compliance, and include side effects (Bangsberg *et al*., 2001; Yazdanpanah, 2004; Hasabi *et al*., 2016; Mehta *et al*., 1997). Furthermore, evolving drug resistance to current regimes is a concern. Consequently, there is a need to develop new therapeutics to combat resistance, increase treatment compliance, and increase accessibility in regions with limited access to healthcare thus preventing infection spread and eradicating HIV-1.

The HIV-1 capsid protein (HIV-1 CA) is integral in the viral life cycle during both early and late stages of infection, but is not targeted by current drug regimens. Compounds such as PF74 and GS-6207 (lencapavir) have demonstrated that HIV-1 CA is a viable drug target (Link *et al*., 2020; Bester *et al*., 2020; Price *et al*., 2014; Bhattacharya *et al*., 2014; Blair *et al*., 2010), targeting a binding pocket between the N-terminal Domain (NTD) and C-terminal Domain (CTD) of adjacent monomers in the capsid hexamer. This binding site is within a cleft present only in the mature hexamer of capsid. It is responsible for binding host factors including CPSF6 and FG-rich peptides of NUP153 (James, 2019; Schaller *et al*., 2011). This binding site presents as a target for drug optimisation or discovery.

The capsid targeting inhibitors, PF74 and GS-6207 (Fig 1.), have been shown to act both during early and late stages of the retroviral lifecycle (Bester *et al*., 2020; Bhattacharya *et al*., 2014). PF74 stabilizes capsid interactions, presumably affecting both capsid uncoating and nuclear entry during the early stages of infection, and capsid maturation during the late stages (Bhattacharya *et al*., 2014). The mechanism of GS-6207 is similar to PF74. The compound GS-6207 readily accelerates capsid formation resulting in aberrant capsid morphology and acts during both early and late stages of the retroviral lifecycle. The principal mode of action is between the reverse transcription and integration steps of the lifecycle (Bester et al., 2020).

**Figure 1.**
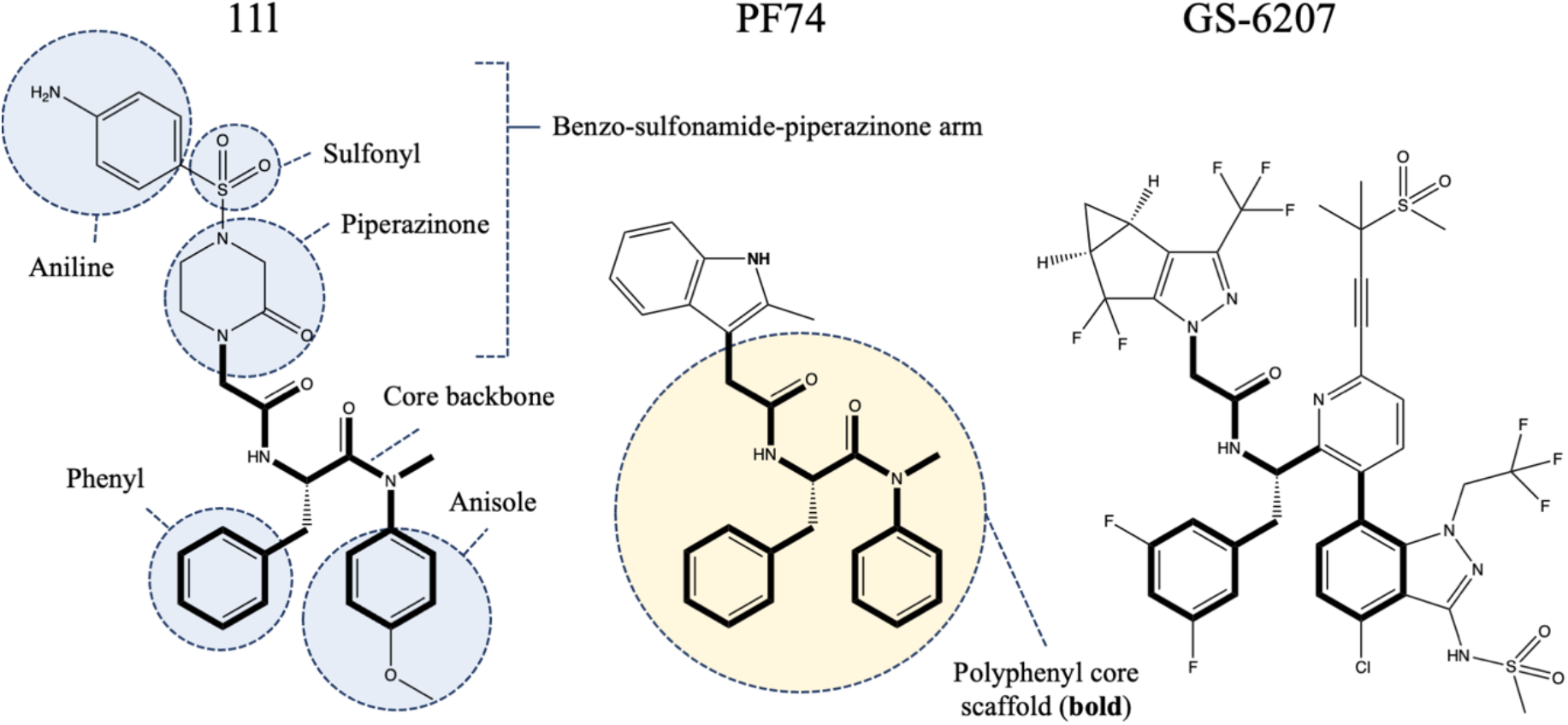
Chemical structure diagrams for capsid inhibitors: 11l, PF74, and GS6207. All compounds share elements of a conserved polyphenyl (bonds shown in bold).

The compound 11l is a third generation antiviral capsid-targeting drug based on polyphenylalanine derivatives of PF74 (Sun *et al*., 2020). In 11l, the indole group has been replaced with a piperazinone, sulfonyl, aniline grouping (Fig. 1). The compound 11l targets both the early and late phases of viral infection, and is 5.8 times more potent at inhibiting infection overall compared to PF74. 11l predominantly acts during the early stages of infection, before reverse transcription, where it is 6.3 times more potent in early-phase restriction compared to PF74 (Sun *et al*., 2020). In contrast to PF74 and GS-6207, 11l appears to accelerate the CA uncoating process and does not affect reverse transcription, indicating a different mode of restriction compared to PF74 and GS-6207.

Here we describe the structure of 11l bound to a disulfide cross-linked hexamer of the HIV-1 capsid protein. The structure shows 11l bound in the same binding pocket as PF-74 and GS-6207. Comparison of the binding pose shows some conformational flexibility and suggests a potential mechanism for the disruption of capsid stability.

## 2. Methods

### 2.1. Preparation of hexameric capsid

Disulfide linked hexamers of the HIV-1 CA protein were formed and purified as previously described (Pornillos *et al*., 2010). HIV-1 CA (A14C/E45C/W184A/M185A) was expressed at scale, harvested, and purified using IMAC and SEC in 10mM Tris/HCl pH 7.8, 150mM NaCl, 20 mM βME. Hexamers were formed via dialysis against 20 mM BME, 150 mM NaCl, 20mM TRIS/HCl pH 8.0, before being transferred to a high-salt solution (20 mM BME, 1M NaCl, 20mM TRIS/HCl pH 8.0) solution overnight at 4 ° C. This promotes the formation of tubular assemblies that mimic the mature capsid lattice seen in HIV-1 virions. These assemblies were then dialysed (1 M NaCl, 20 mM TRIS/HCl pH 8.0) to remove the reducing agent and promote the formation of disulfide bonds between adjacent capsid monomers. In the final dialysis excess salt is removed (150 mM NaCl, 20mM TRIS/HCl pH 8.0) causing tubes to disassemble resulting in disulfide cross-linked hexameric capsid (HIV-1 CA-Hex). Disulfide crosslinked hexameric capsid was purified by SEC using an S200 16/60 column in 10mM Tris/HCl pH 8.0, 150mM NaCl. The assembly was confirmed by non-reducing SDS-page. Purified hexamers were concentrated to 54 mg/ml and stored frozen at −20 ° C.

### 2.2. Crystallisation and X-ray data collection of HIV-1 capsid:11l complex

Initial crystallisation screens were undertaken using the commercially available JCSG, PACT and Morpheus screens (Newman *et al*., 2005; Gorrec, 2009). Crosslinked hexamers were diluted with water to 355 μM (54 mg/mL stock in 150 mM NaCl, 20 mM TRIS/HCl pH 8.0) were mixed 1:1 with 355 μM 11l (from a 50 mM stock in 100% DMSO). After gentle mixing the HIV-1 CA-Hex-11l solution was centrifuged at 13,000 g for 20 minutes. Initial crystals were obtained in 10% PEG8000, 36mM Sodium Malonate, 0.1 M Tris/HCl pH 8.5 but diffracted poorly, consequently were harvested to provide a seed-stock for subsequent seeding and crystallisation.

A single diffracting crystal was obtained in condition D11 of the Morpheus screen (0.12 M Alcohols, 0.1 M Buffer System 3 pH 8.5, 30% v/v Precipitant Mix 3) from a 0.4μL drop consisting of a 0.2μL of 355μM HIV-1 CA-Hex 11l complex supplemented with 10% v/v seed stock and 0.2μL of the reservoir solution. Despite initial precipitation, a single crystal formed after approximately 10 days. The crystal was harvested with a nylon loop and transferred into a cryoprotectant consisting of the crystallisation condition supplemented with 30% ethylene glycol before being frozen in liquid nitrogen.

Data were collected on a Rigaku Synergy-S with a Pilatus 200K detector equipped with an Oxford cryostream device at 110K using a wavelength of 1.54 Å corresponding to the Kα radiation from a copper anode. A single dataset was collected with 0.25° scan width using the Rigaku CrysAlis Pro software.

## 3. Results

Crystals of HIV-1 CA-Hex in complex with the compound 11l were grown in Morpheus screen condition D11 after improvement by iterative rounds of seeding. A single dataset was collected using our in-house X-ray diffractometer. The crystal belonged to the spacegroup P1 with cell dimensions a=90.9 Å, b=91.2 Å, c=116.3 Å, α=87.2°, β=78.8°, γ=60.4°. Inspection of diffraction images identified a non-merohedral twin pathology with the presence of a clear secondary lattice. The two lattices were of similar but not identical dimensions, thus the stronger diffracting lattice, with a higher percentage (48%) of indexed reflections, was selected. Where overlaps occurred intensities were deconvoluted using the CrysAlis Pro software suite. This resulted in a dataset diffracting to 2.5 Å with a CC(1/2) of 0.145 in the outer shell. (See supplementary table 1 for full data collection statistics).

Crystallographic phases were estimated by molecular replacement using an HIV-1 capsid hexamer as a search model (PDB id: 3H47, hexamer generated by symmetry). Two copies of the hexamer were located in the crystal. Each hexamer is present as part of a larger capsid sheet that extends through the crystal, with the two hexamers stacked in adjacent layers (Fig. 2). The two hexamers are offset from one another rather than being stacked perfectly. Inspection of the density for the two hexamers in the unit cell showed clearer density for hexamer I (Chains A-F) compared to Hexamer II (Chains G-L). This is reflected in the average B-factor for each hexamer, with Hexamer II having an average of ∼63.9 Å^2^ compared to Hexamer I ∼32.5 Å^2^. This difference suggests there is some variation in the position of Hexamer II relative to Hexamer I throughout the crystal.

**Figure 2.**
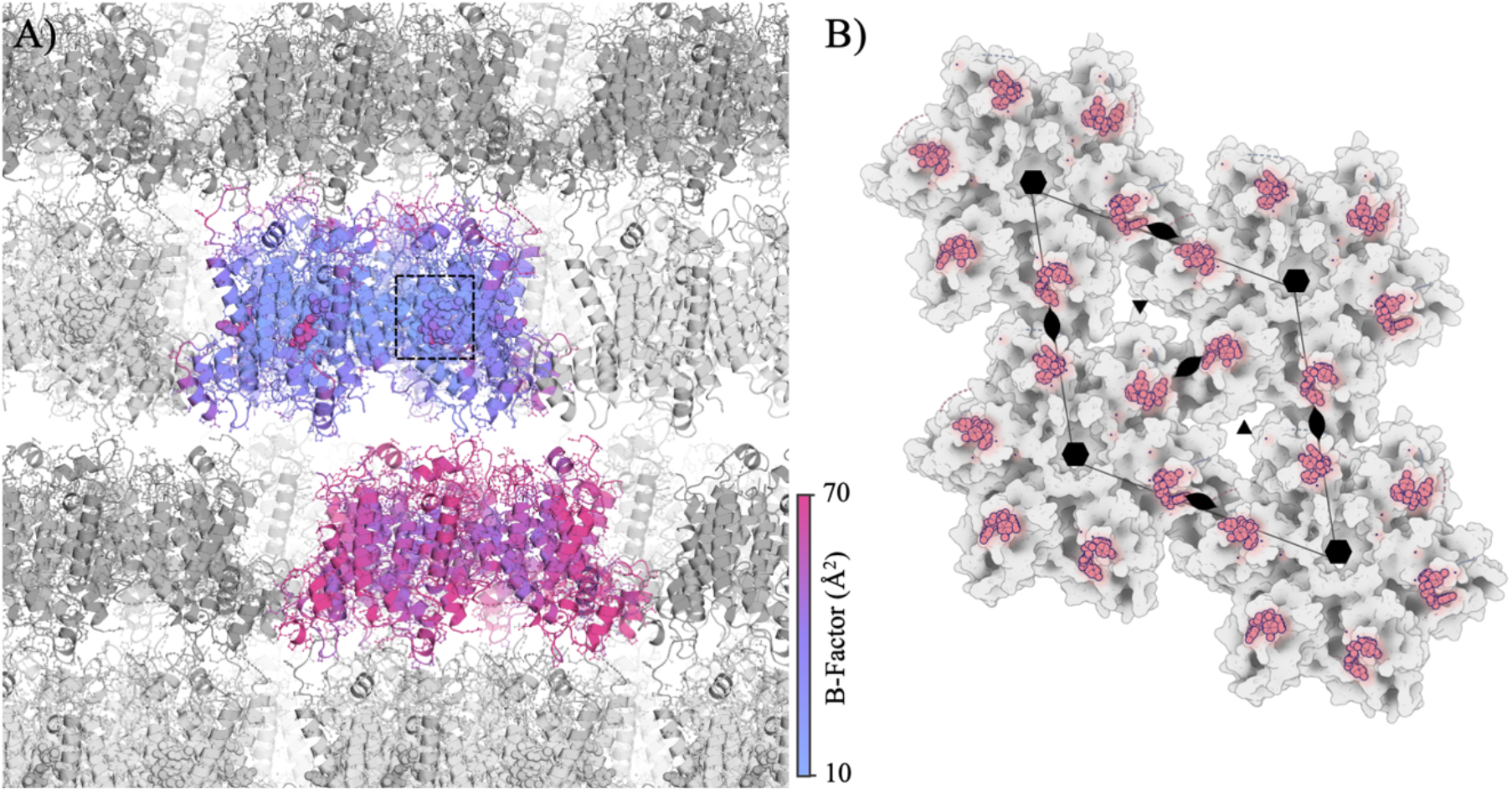
11l binds at the 2-fold HIV-1 capsid interface. **A)** Structure (PDB-ID 8F22) present in unit cell – coloured by B-factor. Shadowed copies show extension of lattice into sheets with alternating offsets. Inset dashed square indicates location and view used in figures 3, and 4. **B)** Structure of hexamer lattice with 11l bound at the 2-fold binding interface between hexamer units.

Iterative cycles of building and refinement resulted in a final Rfact/Rfree of 0.31/0.34. This is considerably higher than other structures at similar resolution, and is likely due to the loss of data quality from the deconvolution of non-merohedral twins. The measure of each intensity is affected by its percentage of overlap with a twinned reflection, a greater over-lap results in poorer intensity estimates during deconvolution, or lost data if the overlap is >95%.

Inspection of the electron density maps revealed density corresponding to the bound 11l was in the pocket that is known to bind PF74 and GS6207. This pocket is found within the cleft formed between the NTD domains of tiling hexamers and is situated between the NTD and CTD of adjacent monomers within the hexamer, bounded by helices α3, α4, and α5 of the NTD (Fig. 3B). The sidechain from Arg173 mediates hydrogen bonds with the mainchain carboxyl groups of Asn57 and Val59 from the NTD of the adjacent monomer and directly contributes to the wall of the pocket with Lys 70 and Gln67 bounding the other side. The pocket is adjacent to the loop between α8 and α9 of the neighbouring monomer within the hexamer which mediates the 2-fold interactions to other hexamers of the lattice. In our structure this loop is poorly ordered and not visible in the density. The 11l compound occupies this pocket and forms interactions with both the NTD chain, and the CTD of the neighbouring monomer within the hexamer.

**Figure 3.**
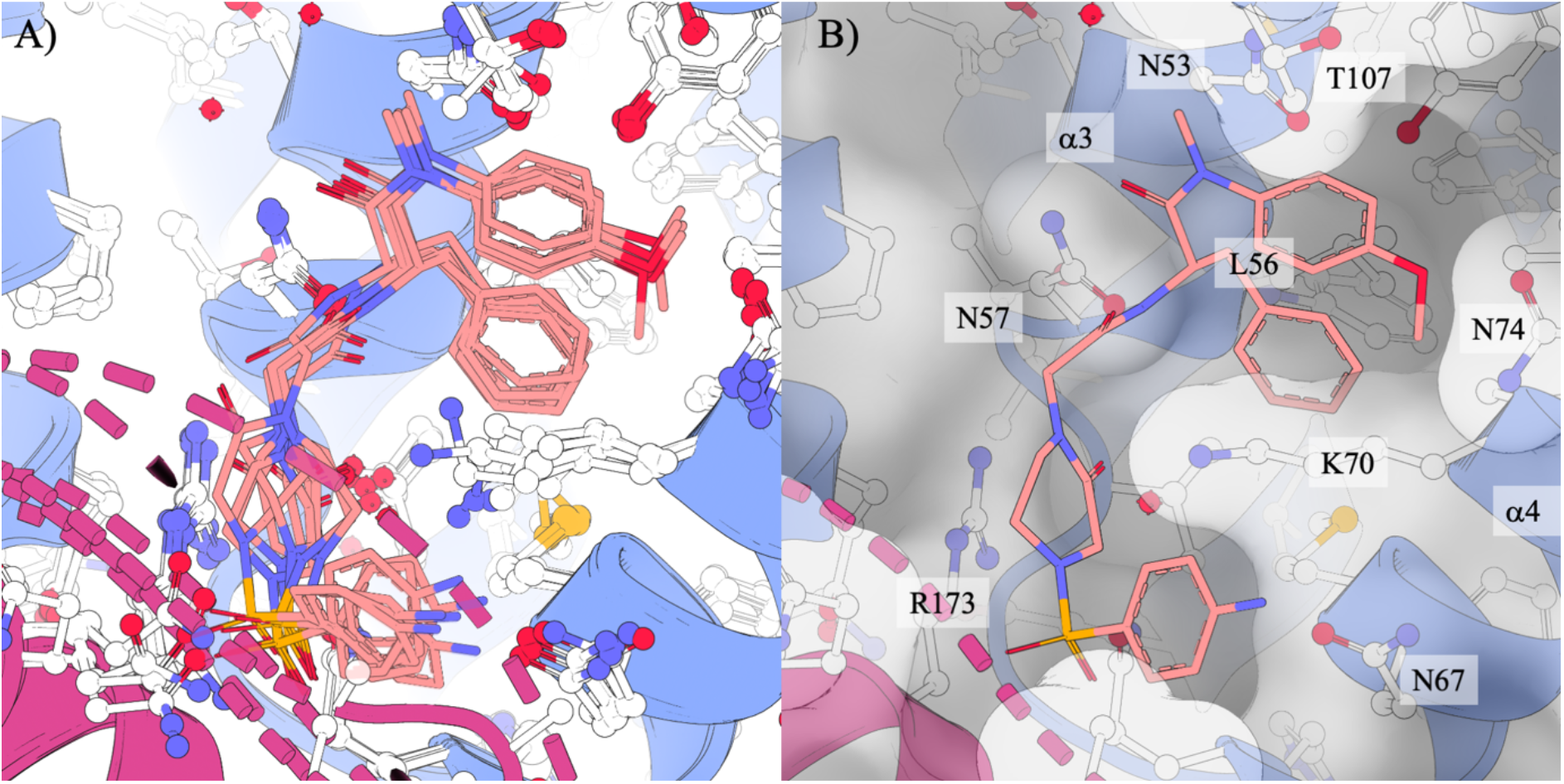
Structures of 11l bound to capsid binding site A) overlay of all 6 binding sites from chains A, B, C, D, E, and F within the model. The core scaffold is consistent, confirmational variability is seen across the benzo-sulfonamide-piperazinone arm and anisole group. B) Close view of a single bound 11l compound (Chain D).

Inspection of this pocket showed clear density for a ligand present in the six sites of Hexamer I only. The equivalent sites in hexamer II shows signs of the presence of the compound however the inflated B-factors make the density difficult to interpret and consequently we have not modelled the 11l compound in Hexamer II. The 11l compound is present in hexamer I with a topology similar to the binding mode of PF74. The density allows all components of the compound to be placed at full occupancy with B-factors equivalent to the surrounding protein chain.

The aromatic groups of the anisole and phenyl groups within the polyphenyl core of 11l occupy the same binding pockets as the R1 and R2 groups of PF74 (Fig. 4). The anisole group occupies the space between Lys70, Thr107, and terminates variably with the methoxy end orientated towards or away from Asn74 when comparing across chains. The bridging backbone atoms between the anisole and phenyl group consists of a methyl group, protruding from the nitrogen atom connecting the anisole, and a carboxyl group before starting the phenylalanine-like central structure. This methyl and carboxyl group are planar and orientated away from the binding pocket. The methyl group points towards the main chain carboxyl of Gly106, however it is too distant to contribute to any major interactions with any of the mainchain backbone. The carboxyl group of 11l orients and hydrogen bonds with the ND2 atom of Asn53. A second nitrogen atom in the backbone scaffold of 11l forms a second hydrogen bond with the OD1 atom of Asn53. The phenyl moiety occupies a deep hydrophobic pocket bounded by the residues Leu56, Leu69, and Ile72. The sidechain of Lys70 sits above the phenyl group, resulting in the phenyl being entirely buried. The polyphenyl core in this regard is identical in its binding modality to the equivalent structure in PF74 suggesting it is a major driver of the binding pose.

**Figure 4.**
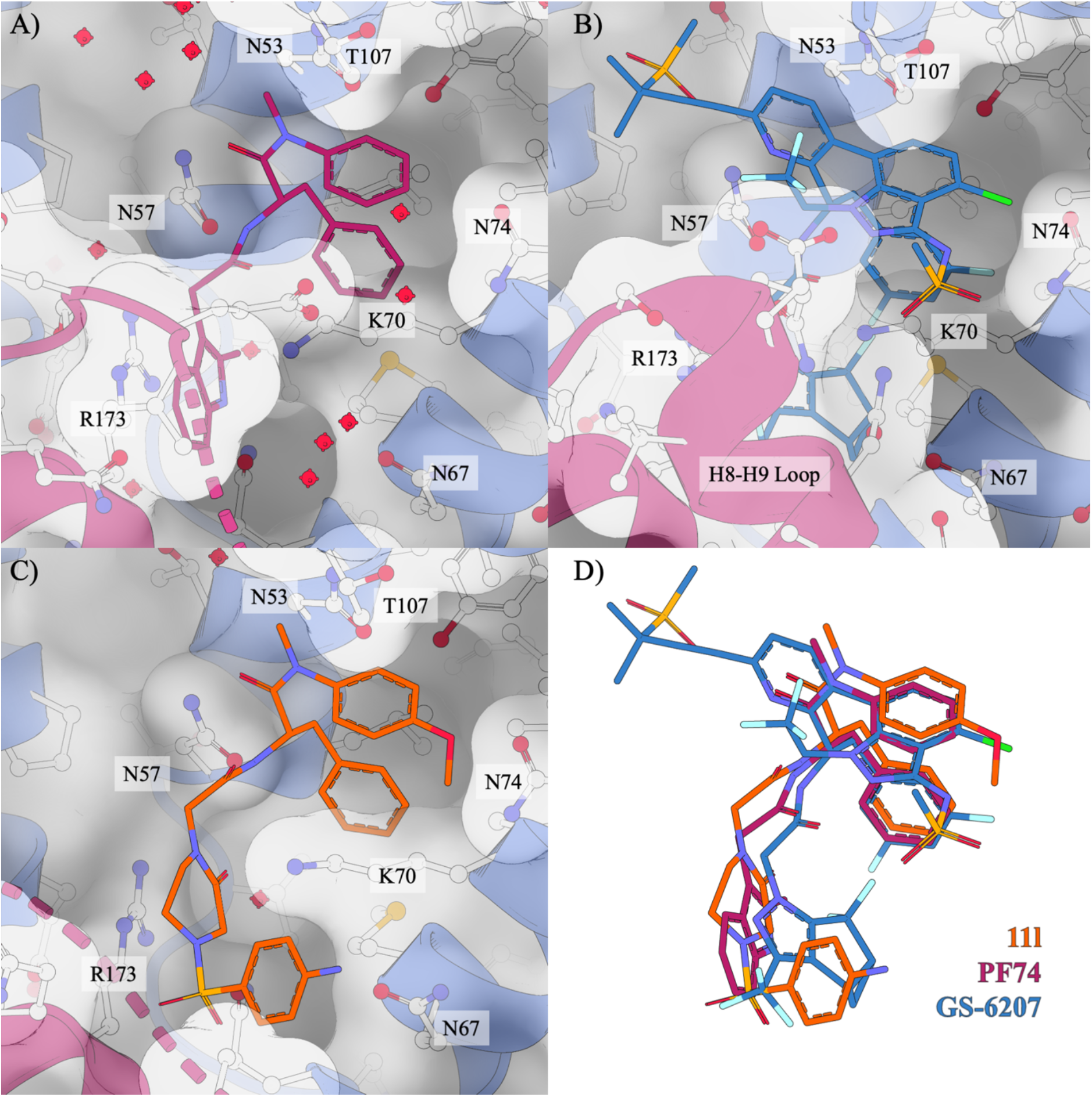
Comparison of 11l, PF74, and GS6207 binding. All compounds exhibit a shared mode of binding. Only GS-6207 fully stabilises the H9-H9 loop. Only a portion of the loop is visible when PF74 is bound. The loop is missing when 11l is bound. A) PF74, B) GS-6207, C) 11l, D) overlay of three compounds.

The piperazinone group is located in the equivalent position to the indole group of PF74. Comparing across chains it displays a high degree of mobility. The piperazinone ring is a saturated ring system and can display conformational flexibility providing an explanation for the disorder seen. Additionally, it is able to rotate along its axis, positioning the ketone group into two predominant orientations: interacting with the Nζ atom of Lys70, or the Nε atom of Arg173 from the adjacent chain. In chain C and E, a well-ordered water molecule is present co-ordinating the ketone, as well as favourable polar interactions with the sidechains of Gln63, and Arg173 from the adjacent monomer. In Chain A, 1ll is modelled with the piperazinone ketone group directed away from the binding site. All other chains show evidence that the preferred orientation is with the ketone group oriented towards the CTD Arg173 Nε atom from an adjacent chain, the Lys70 Nζ atom, or the nearby water molecule.

Despite disorder and variability in the piperazinone moiety position, the sulfonyl group is present in the same position for all 11l compounds modelled. The disorder of the adjacent loop (between CTD helix 8 and 9) in the capsid makes interactions in this region difficult to assess. No hydrogen bonds to neighbouring amino acids are identified. Instead, the sulfonyl is located in a region of general positive charge making van der walls and general electrostatic interactions with the binding pocket.

The final moiety of 11l is the aniline group, which for all instances of 11l is positioned towards space occupied by the loop between CTD helices 8 and 9 in unliganded capsid. The planar ring of the aniline shows rotational freedom with a different orientation in each chain.

## 4. Discussion

The capsid is crucial to the lifecycle of the retrovirus, it mediates active transport through the cytoplasm, transport through the nuclear pore complex, evasion of innate immune sensors, and the timely uncoating releasing the viral genome for integration and the maturation of newly formed virions (James, 2019; Campbell & Hope, 2015; Carnes *et al*., 2018; Zila *et al*., 2020). Interfering with any of these functions has the potential to compromise infection at multiple points of the life-cycle, making the capsid an important target for the development of antiretrovirals. Future capsid binding drugs have potential to make a significant impact to treatment and trajectory of the HIV-1 pandemic.

The compound 11l is derived from PF74 sharing a polyphenyl core that mimics the binding of the FG repeats present in the nuclear import protein NUP153. The structure presented here shows the core scaffold of 11l binds identically to its derivative PF74. The phenyl and anisole groups occupy the same hydrophobic pockets mediated in part by Leucine (Leu56, Leu69) and Isoleucine (Ile73) residues, whilst the phenyl group is effectively buried by the long sidechain of Lysine (Lys70). The only major difference between the phenyl core of 11l and PF74 is the methoxy group within the anisole moiety present only on 11l, which is in one of two preferred orientations.

More recently the PF74 derivative GS-6207 (lencapavir) has obtained regulatory approval from the European Commission for the treatment of multi-drug resistant HIV-1. Comparison of 11l to GS-6207 demonstrates that despite significant deviations from the core, the binding topology of GS-6207, PF74, and 11l all share the same binding mode, with the two core phenyl or phenyl derivatives occupying the hydrophobic pockets between Lys70 and Leu56, and Lys70 and Thr107. To note, GS-6207 contains an indole-like moiety, similar to PF74, which occupies the equivalent position of the indole of PF74. In this way, 11l is unique when compared to GS-6207 and PF74 for the extended reach of the aniline moiety. When comparing structures of 11l to PF74 and GS-6207, the region with the largest changes are residues 175-190, i.e., the region starting from the terminus of helix 8, through to helix 9, that forms a key part of the hexamer-hexamer interface in the extended capsid lattice. This loop can be poorly ordered in the structures of HIV-1 CA, depending on the construct used (Bhattacharya *et al*., 2014; Pornillos *et al*., 2009; Gres *et al*., 2015), with poorly defined electron density that is generally not modelled reflecting an area that exhibits conformational variability. Binding of PF74 increases the order of residues 176-180, making them visible in the crystal structure. Binding of GS-6207 orders the entire loop, forms favourable interactions with the compound. In the structure of 11l, this loop is partially ordered with residues 175-178, and 184-190 visible. Residues 179-183 are not visible in the map suggesting this as a possible area for future improvements. It must be noted that two mutations used in this study (and the studies of PF-74 and GS-6207) W184A/M185A are located in this region and may contribute to the disorder. There are also differences in crystal packing that may be responsible for differences in the stability of the loop. This loop plays an important role in the 2-fold symmetry Ctd-Ctd interactions that mediate formation of the extended capsid lattice.

### 4.1. 11l proposed mechanism of action

Both PF74 and GS-6207 are thought to act by stabilising the capsid architecture, increasing the rate of capsid assembly *in vivo*. This affects maturation in the late phases of infection. Additionally, competition between these compounds and the nuclear import factors CPSF6 and NUP153 are thought to disrupt early stages of the viral lifecycle (Matreyek *et al*., 2013; Schaller *et al*., 2011; Bhattacharya *et al*., 2014). In contrast 11l has been shown to slow assembly of capsid protein in capsid assembly assays (Sun *et al*., 2020). This is further supported by the destructive nature 11l had on unliganded crystals during soaking experiments when compared to PF74 that can be readily soaked into crystals of HIV-1 CA (data not shown). This would support the hypothesis that 11l is affecting capsid assembly, and potentially mediating disassembly, at a level that is not congruent with the formation of lattices, or mature capsid icosahedra.

The binding pose of 11l seen in our structure provides some suggestions of how this might be occurring. The polyphenyl core functions by tightly binding in the FG pocket of the hexamer effectively targeting and locking the ligand to the mature capsid, while the piperazinone, sulfonyl aniline arm is less well positioned and disrupts positioning of the helix 8-helix 9 loop of the capsid thus destabilising the capsid core lattice. This would provide a reasoning for the more potent early phase restriction profile of 11l compared to PF74.

### 4.2. Pathways for future development

Structural optimisation of the 11l compound may therefore benefit from strengthening the poly-phenyl core interaction with the NTD, whilst also maintaining steric hindrance and unfavourable interactions to the adjacent monomer CTD. Importantly, one could expect the steric hindrance properties of the benzosulfonamide-piparizinone group may not be improved, without first ensuring the polyphenyl cores binding affinity is strong enough to keep the inhibitor in the binding pocket. Assuming this interpretation, the interaction between the poly-phenyl core region of 11l and the HIV-1 CA assembly should be focused for increased affinity, before further screening of motifs in the sulfonamide aniline arm. GS-6207 provides insights into how this may be achieved: fluoridation of the phenyl group, and extension of the polyphenyl-core back-bone. Furthermore, unmodelled solvent density between residues Thr107 and Asn74 suggest that there is room to exploit additional interactions with the protein. The methoxy end of the anisole group may be extended to displace this solvent, reducing methoxy disorder, and increasing the poly-phenyl core binding affinity.

**Table 1.** Crystallographic data collection and refinements statistics.

Crystallographic data and refinement statistics for PDB 8F22
|  |  |
| --- | --- |
| Wavelength | 1.543 |
| Resolution range | 15.89 - 2.5 (2.589 - 2.5) |
| Space group | P 1 |
| Unit cell | 91.0 91.2 116.3 87.2 78.8 60.4 |
| Total reflections | 209105 (21512) |
| Unique reflections | 106374 (10897) |
| Multiplicity | 2.0 (2.0) |
| Completeness (%) | 96.64 (99.02) |
| Mean I/sigma(I) | 4.13 (2.41) |
| Wilson B-factor | 20.74 |
| R-merge | 0.152 (0.288) |
| R-meas | 0.216 (0.407) |
| R-pim | 0.153 (0.288) |
| CC1/2 | 0.969 (0.145) |
| CC* | 0.992 (0.504) |
| Reflections used in refinement | 106368 (10897) |
| Reflections used for R-free | 5353 (581) |
| R-work | 0.313 (0.366) |
| R-free | 0.345 (0.387) |
| CC(work) | 0.886 (0.218) |
| CC(free) | 0.837 (0.185) |
| Number of non-hydrogen atoms | 17395 |
| macromolecules | 17130 |
| ligands | 246 |
| solvent | 19 |
| Protein residues | 2235 |
| RMS(bonds) | 0.29 |
| RMS(angles) | 4.34 |
| Ramachandran favoured (%) | 98.68 |
| Ramachandran allowed (%) | 1.32 |
| Ramachandran outliers (%) | 0 |
| Rotamer outliers (%) | 2.07 |
| Clash-score | 9.67 |
| Average B-factor | 45.8 |
| macromolecules | 45.8 |
| hexamer A-F | 32.5 |
| hexamer G-L | 63.9 |
| ligands | 42.4 |
| solvent | 20.1 |
| Number of TLS groups | 85 |

Data and refinement statistics (high resolution shell in parentheses).

